# Unveiling the Cellular and Molecular Mechanisms of Diabetic Retinopathy with Human Retinal Organoids

**DOI:** 10.1101/2025.02.24.639921

**Authors:** Lada Polešovská, Simona Trmačová, Canan Celiker, Eva Hrubá, Francisco Molina Gambin, Veronika Matušková, Eleni Beli, Tomáš Bárta

## Abstract

Diabetic retinopathy (DR) is a leading cause of vision impairment worldwide, driven by chronic hyperglycaemia and its complex metabolic consequences. While animal models have been widely used to study DR, they often fail to replicate the physiology of human retina. To address this limitation, we employed human retinal organoids as a model to study the effects of hyperglycaemia across various stages of retinal differentiation. Early-stage organoids demonstrated resilience to high glucose levels, maintaining normal morphology, viability, and gene expression. However, advanced-stage organoids displayed significant disruptions, including the downregulation of outer segment-specific genes, which impaired photoreceptor maturation, and a noticeable shortening of photoreceptor outer segments. Transcriptomic analysis revealed substantial changes in pathways related vision including G protein-coupled receptor signalling pathway, response to light stimulus, and visual perception. While photoreceptors were particularly vulnerable, other retinal cell types, including bipolar cells, ganglion cells, and Müller glia, showed greater resilience. Additionally, glial activation, evidenced by increased expression of astrocyte markers, suggested an adaptive response to hyperglycaemia. To validate our findings, we compared our dataset with publicly available transcriptomic datasets from human retinas with DR, confirming key overlaps in pathways related to photoreceptor dysfunction, gliogenesis, and oxidative stress responses. These results establish human retinal organoids as an effective and relevant model for studying the molecular mechanisms of neurodegeneration associated with DR progression.

## Introduction

Diabetic retinopathy (DR) is one of the most prevalent complications of diabetes mellitus and stands as a leading cause of vision loss among adults worldwide. The global prevalence of DR was 103 million in 2020 and is expected to reach 160.5 million by 2045 ^1^. DR results from chronic hyperglycaemia, which triggers metabolic cascades that progressively damage the retina, leading to visual impairment and blindness. Given the increasing incidence of diabetes globally, DR is a critical focus within both ophthalmologic and systemic diabetic care.

Traditionally, DR has been characterized as a retinal microvascular disorder leading to vascular alterations such as capillary leakage, occlusion, and subsequent ischemia ^2^. However, this vascular-centred view has been recently challenged by accumulating evidence suggesting that retinal neurodegeneration may precede microvascular pathology, contributing to the initiation and progression of DR ^3^. Studies indicate that retinal ganglion cells, photoreceptors, and other retinal neurons may undergo early, diabetes-induced degenerative changes, setting the stage for the later vascular complications ^4^.

Additionally, diabetes mellitus during pregnancy introduces an often-overlooked complication that can impact both the mother and the developing foetus. Maternal hyperglycaemia exposes the foetal retina to elevated glucose levels, potentially disrupting early retinal development and increasing the risk of long-term visual impairment ^5,6^. Despite this, we lack suitable models to study how hyperglycaemia affects the developing retina. Establishing such models is essential for understanding gestational DR and its impact on foetal retinal health.

Current DR research relies heavily on animal models to elucidate disease mechanisms and assess potential therapies. However, animal models present limitations, as they often fail to fully replicate human retinal physiology and pathology, especially with respect to the neurodegenerative components of DR. These challenges highlight the need for human-based *in vitro* models that can accurately recapitulate the complex cellular architecture and functional attributes of the human retina.

Here, we utilized human retinal organoids as a versatile and scalable model to investigate the effects of hyperglycaemia on retinal development and function, simulating the conditions of gestational DR. Our findings reveal that early-stage retinal organoids exhibit remarkable resilience to hyperglycaemia, with no significant morphological, viability, or molecular changes. However, as the organoids progress through differentiation, hyperglycaemia induces notable molecular disruptions, including the downregulation of photoreceptor-specific genes, an enhanced oxidative stress response, and reduced proliferative activity. These changes indicate the vulnerability of photoreceptors to chronic glucose exposure while demonstrating resilience in other retinal cell types including bipolar cells, ganglion cells, and Müller glia. Our transcriptomic analysis reveals distinct gene expression profiles and enriched pathways disrupted by hyperglycaemia, including phototransduction, sensory perception, and G protein-coupled receptor signalling pathway. Furthermore, we observed glial activation, marked by upregulation of astrocytic markers, indicative of a reactive gliosis response in hyperglycaemic conditions.

In this study, we establish human retinal organoids as a powerful model for investigating the cellular and molecular mechanisms of DR, offering a human-specific system to uncover the temporal and cell-specific impacts of hyperglycaemia and explore potential therapeutic targets.

## Results

### High glucose levels do not affect retinal organoids during the early stage of differentiation process

We investigated the effects of hyperglycaemia on different stages of retinal organoid differentiation. Retinal organoids were differentiated for 30 days, 90 days, and 150 days (D30, D90, D150), followed by 28 days of treatment with either L-glucose (Osmotic control, consisting of the 17.5 mM D-glucose already present in the culture medium plus an additional 7.5 mM L-glucose), D-glucose (Hyperglycaemia, consisting of the 17.5 mM D-glucose already present in the culture medium plus an additional 7.5 mM D-glucose, total 25 mM D-glucose), or the original organoid medium (Control, 17.5 mM D-glucose) (**Fig. 1A**). It is important to note that while 17.5 mM D-glucose already represents a hyperglycaemic condition compared to physiological levels, this concentration is commonly used in organoid differentiation protocols as a standard baseline to ensure robust development and survival of organoids *in vitro* ^7,8^. Therefore, it provides a well-accepted framework for studying the additional effects of higher glucose concentrations. Furthermore, the inclusion of an osmotic control (L-glucose) helps to separate the specific effects of glucose metabolism from osmotic stress. This experimental design ensures that the observed differences are attributable to hyperglycaemia-induced molecular and cellular responses, rather than variability in baseline culture conditions. Additionally, we attempted to culture retinal organoids in 5.5 mM D-glucose to mimic physiological glucose levels. However, these organoids did not survive beyond seven days, as they disintegrated rapidly (**Fig. S1A**).

**Figure 1:**
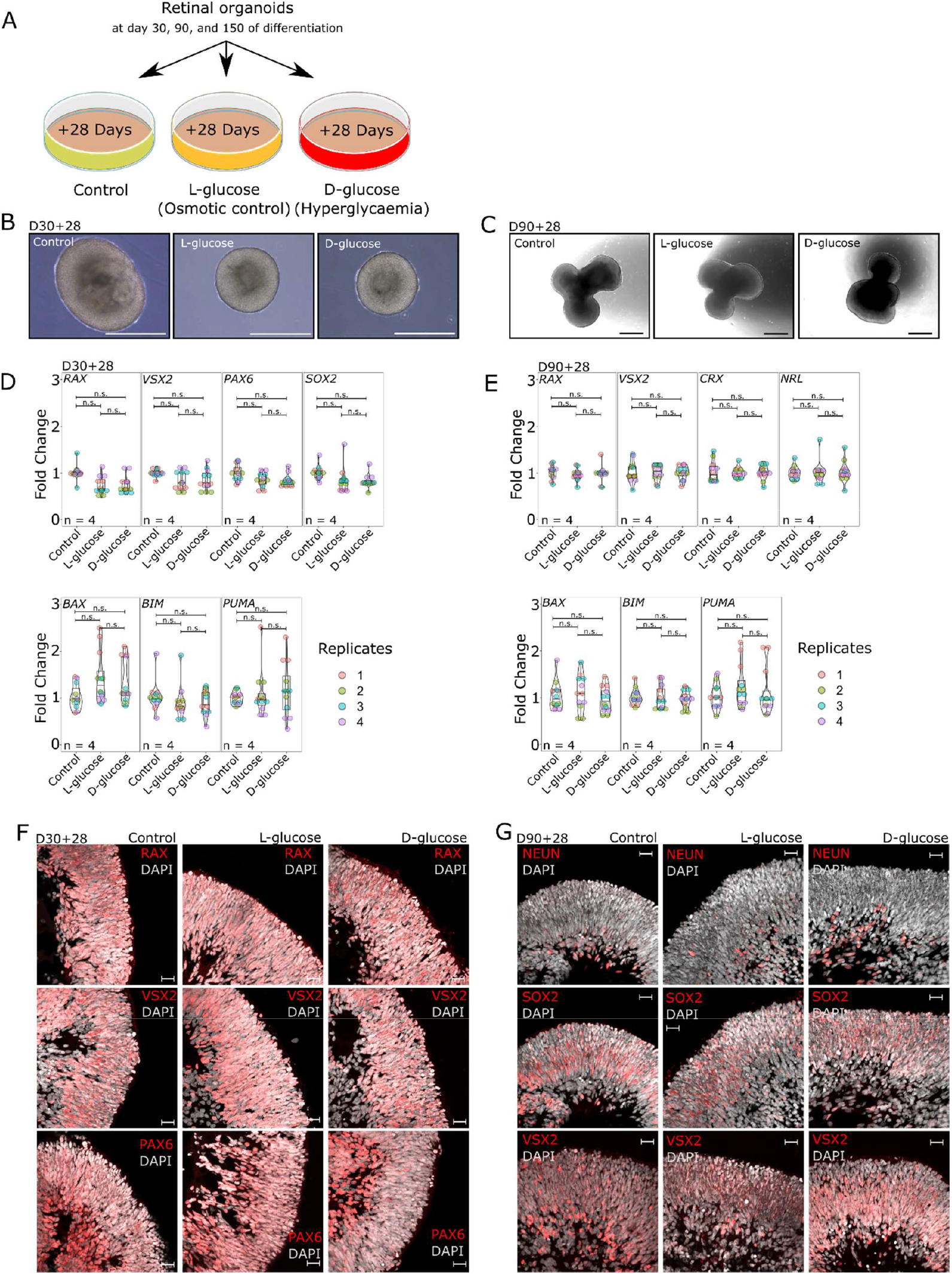
High glucose levels do not affect retinal organoids during early differentiation. **A)** Experimental design of the whole study. Retinal organoids were differentiated for 30, 90, and 150 days (D30, D90, D150) and then treated for 28 days with either Control medium (17.5 mM D-glucose), L-glucose (osmotic control, 17.5 mM D-glucose + 7.5 mM L-glucose), or D-glucose (hyperglycaemia, 25 mM D-glucose). **B)** Representative brightfield images of retinal organoids at D30+28 for Control, L-glucose, and D-glucose treatment groups, demonstrating similar morphology and no obvious structural abnormalities. Scale bars: 500 µm. **C)** Representative brightfield images of retinal organoids at D90+28, showing characteristic neuroepithelial morphology across all conditions. Scale bars: 200 µm. **D, E)** RT-qPCR analysis of gene expression. Results were obtained from retinal organoids derived from two independent cell lines. Individual data points are shown. **D)** Early retinal and neuronal markers (RAX, VSX2, PAX6, SOX2) and apoptosis-related genes (BAX, BIM, PUMA) at D30+28 and **E)** retinal development markers (RAX, VSX2, CRX, NRL) and apoptosis-related genes (BAX, BIM, PUMA) at D90+28 were analysed. **F, G)** Indirect immunofluorescence staining. **F)** Representative images of early retinal markers (RAX, VSX2, PAX6) at D30+28. **G)** Representative images of retinal and neuronal markers (NEUN, SOX2, VSX2) at D90+28. All markers show normal expression and spatial localization across conditions. DAPI marks nuclei. Scale bars: 20 µm.

Our data indicate that the early stages (D30+28, D90+28) of retinal organoid differentiation are not significantly affected by hyperglycaemic conditions. Retinal organoids cultured under high glucose conditions showed no signs of increased cell death and exhibited normal morphologies comparable to control groups. Across all conditions, the organoids successfully formed neuroepithelial structures with characteristic organization for this stage of differentiation (**Fig. 1B, C**).

To further investigate the potential impact of hyperglycaemia on early retinal development, we conducted RT-qPCR analysis to quantify the expression levels of retinal and neuronal markers. Specifically, *RAX, VSX2, PAX6*, and *SOX2* were analysed at D30+28, while *RAX, VSX2, CRX*, and *NRL* were analysed at D90+28. As shown in **Figures 1D and 1E**, there were no statistically significant changes in the expression levels of these genes among the control, L-glucose (osmotic control), and D-glucose (hyperglycaemia) groups. This suggests that the high glucose environment does not adversely influence the expression of these critical developmental genes. In addition to retinal markers, we assessed the expression levels of apoptosis-related genes (*BAX, BIM*, and *PUMA*) using RT-qPCR. Our results showed no significant differences in the expression of these genes across the treatment groups, suggesting that hyperglycaemia does not induce apoptosis in retinal organoids during the early stages of differentiation (**Fig. 1D, E**).

Finally, we examined the localization and distribution of key retinal and neuronal transcription factors including RAX, VSX2, and PAX6 at D30+28, and NEUN, SOX2, and VSX2 at D90+28 in the retinal organoids. Consistent with our RT-qPCR findings, immunofluorescence analysis revealed no detectable abnormalities in the spatial expression patterns of these markers across the different treatment groups. Representative images in **Figures 1F and 1G** demonstrate that RAX, VSX2, PAX6, NEUN, and SOX2 are expressed within the neuroepithelial structures of the retinal organoids, with no visible differences between control and hyperglycaemic conditions.

These results suggest that high glucose conditions do not affect the early differentiation, structural development, or cell survival of retinal organoids. The preservation of typical morphology, gene expression levels, apoptosis gene regulation, and protein localization under hyperglycaemic conditions indicates that early retinal differentiation remains resilient to the metabolic challenges associated with elevated glucose levels.

### High glucose levels impact photoreceptor gene expression in late-stage retinal organoids

Building on our findings from earlier differentiation stages, where hyperglycaemic conditions did not significantly impact retinal organoid morphology, gene expression, or protein expression and localization, we next investigated whether these effects persisted or emerged during the later stages of development. At D150+28, retinal organoids are at an advanced stage of differentiation, with expected progression of photoreceptor maturation.

Brightfield microscopy images of the organoids **(Fig. 2A)** revealed no morphological differences across the treatment groups (Control, L-glucose, and D-glucose), like the observations in earlier stages of differentiation. However, in contrast to the earlier stages, significant differences in key retinal-specific gene expression emerged at D150+28.

**Figure 2:**
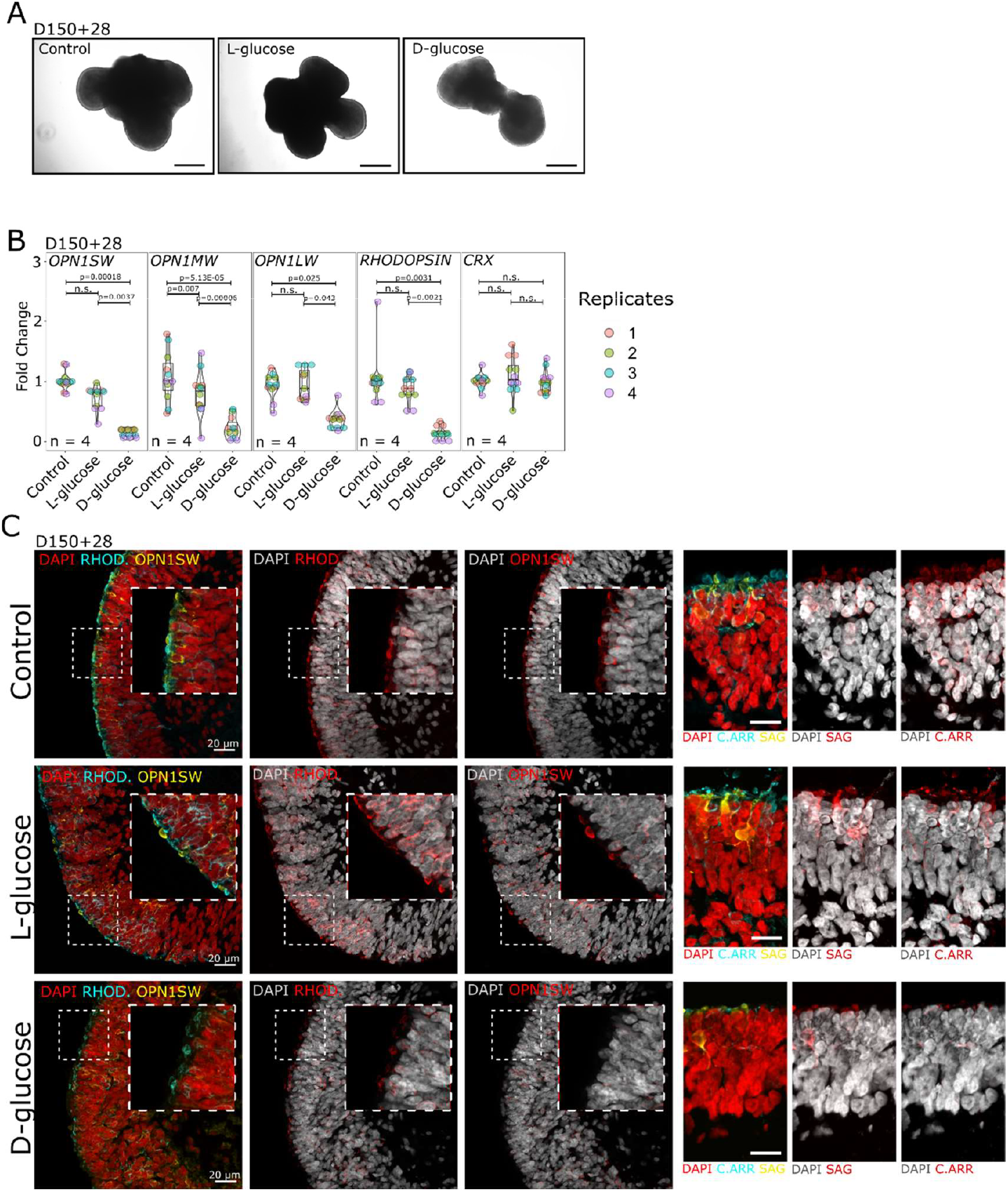
High glucose levels impact photoreceptor gene expression in late-stage retinal organoids. **A)** Brightfield microscopy images of retinal organoids in Control, L-glucose, and D-glucose conditions. No morphological differences were observed between groups. Scale bar: 100 µm. **B)** RT-qPCR analysis of photoreceptor-specific genes OPN1SW, OPN1MW, OPN1LW, and RHODOPSIN shows significant downregulation in the D-glucose group compared to Control and L-glucose groups. CRX expression remains unchanged across all conditions. Results were obtained from retinal organoids derived from two independent cell lines. Individual data points are shown. **C)** Immunofluorescence staining of retinal organoids for photoreceptor markers. Left panel: OPN1SW and RHODOPSIN staining reveal reduced expression in the D-glucose group compared to Control and L-glucose groups. Right panel: C.ARR (Cone arrestin, AAR3) and SAG (Rod arrestin) also exhibit reduced expression in the D-glucose group. DAPI marks nuclei. Insets show magnified regions of interest. Scale bars: 20 µm.

RT-qPCR analysis was performed to measure the expression levels of photoreceptor-specific genes, including *OPN1SW* (short-wavelength-sensitive opsin), *OPN1MW* (medium-wavelength-sensitive opsin), *OPN1LW* (long-wavelength-sensitive opsin), and *RHODOPSIN*. The results showed a significant downregulation of expression of all four genes in the D-glucose group compared to the controls (**Fig. 2B**), suggesting that D-glucose treatment negatively impacts the expression of critical photoreceptor markers. Notably, L-glucose also led to a significant downregulation of *OPN1MW*, despite being an osmotic control. This indicates that osmotic stress alone can influence the expression of certain genes, potentially through secondary effects on cellular homeostasis or metabolic pathways. However, *OPN1SW, OPN1LW* and *RHODOPSIN* remained unaffected in L-glucose, suggesting that the effects of D-glucose on these genes are likely due to hyperglycaemia rather than osmotic stress. Interestingly, *CRX*, a key transcription factor involved in photoreceptor development, remained unchanged across all conditions (**Fig. 2B**).

Immunofluorescence staining revealed reduced outer segment (OS) structures and expression of photoreceptor markers in the D-glucose group compared to the control and L-glucose groups (**Fig. 2C**). Both OPN1SW and RHODOPSIN showed decreased levels, consistent with the RT-qPCR results. Additionally, cone arrestin (C.ARR, ARR3) and SAG (S-antigen or arrestin-1) were also downregulated, indicating a broader disruption in photoreceptor differentiation under D-glucose conditions.

Having observed the downregulation of photoreceptor-specific genes in response to hyperglycaemia, we next investigated whether this might be associated with apoptosis, particularly in photoreceptors. We performed RT-qPCR analysis of key pro-apoptotic genes, including *BAX* and *PUMA* (**Fig. 3A**). However, no significant differences were observed between the control, L-glucose, and D-glucose groups, suggesting that the transcriptional regulation of apoptosis-related genes remains unchanged across conditions. Additionally, we performed a more comprehensive analysis of apoptosis-related proteins using protein arrays (**Fig. 3B, Fig. S1B, C, D**). This analysis also revealed no upregulation or activation of apoptosis-related markers, supporting the notion that apoptosis is not broadly induced at the protein level under D-glucose treatment.

**Figure 3.**
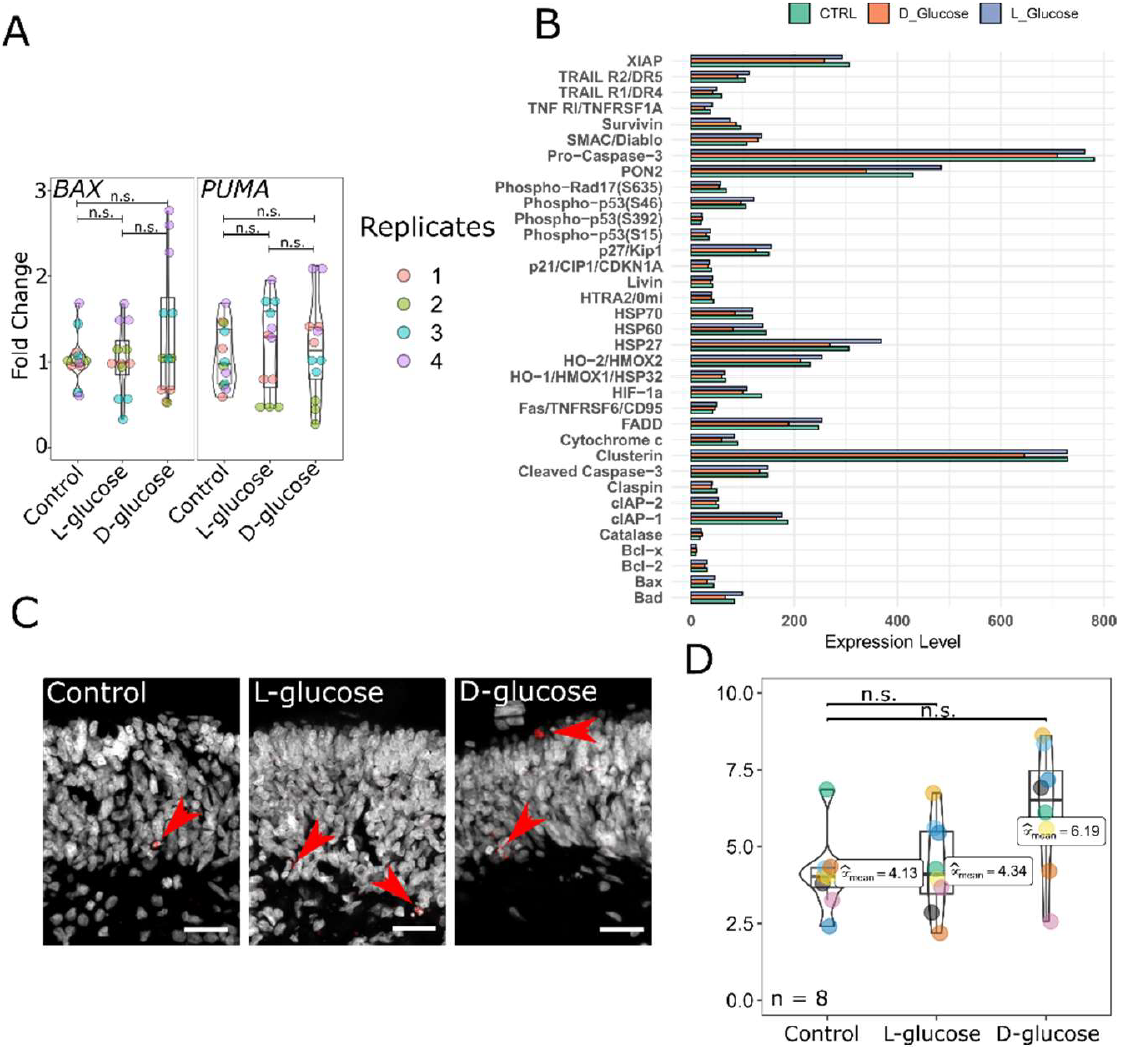
Analysis of Apoptosis-Related Responses in Late-Stage Retinal Organoids Exposed to Hyperglycaemia. **A)** RT-qPCR analysis of pro-apoptotic genes (BAX and PUMA) in retinal organoids after D-glucose, L-glucose, or control treatment at D150+28. No significant changes in expression levels were observed across conditions. Results were obtained from retinal organoids derived from two independent cell lines. Individual data points are shown. **B)** Quantification of apoptosis-related proteins using a protein array. **C)** Representative immunofluorescence images of retinal organoids stained for cleaved caspase-3, a marker of apoptosis. Red arrowheads indicate cleaved caspase-3-positive cells. Scale bar = 50 µm. **D)** Quantification of cleaved caspase-3-positive cells in retinal organoids.

Recognizing that bulk approaches may obscure cell-specific responses, we turned to immunofluorescence staining for cleaved caspase-3, a marker of apoptosis, to localize apoptotic activity within the organoids (**Fig. 3C**). Quantification of cleaved caspase-3-positive cells (**Fig. 3D**) showed an elevated (∼1.5 fold increase), but not statistically significant, number of apoptotic cells in the D-glucose group compared to the controls. These findings suggest that while D-glucose treatment does not induce widespread apoptosis in the organoids, there may be a localized or cell-specific apoptotic response, potentially affecting photoreceptors more prominently.

### Transcriptomic Analysis Revealed Profound Alterations in Photoreceptor-Specific Gene Expression

To gain a comprehensive understanding of the molecular changes induced by hyperglycaemia, we performed RNA sequencing on retinal organoids treated with D-glucose (D150+28) compared to controls. Principal component analysis revealed distinct transcriptomic profiles between the control and D-glucose-treated organoids (**Fig. 4A**). Control replicates formed a tightly clustered group, while D-glucose replicates grouped separately, indicating that hyperglycaemic treatment induces widespread and reproducible changes in gene expression patterns within retinal organoids.

**Figure 4.**
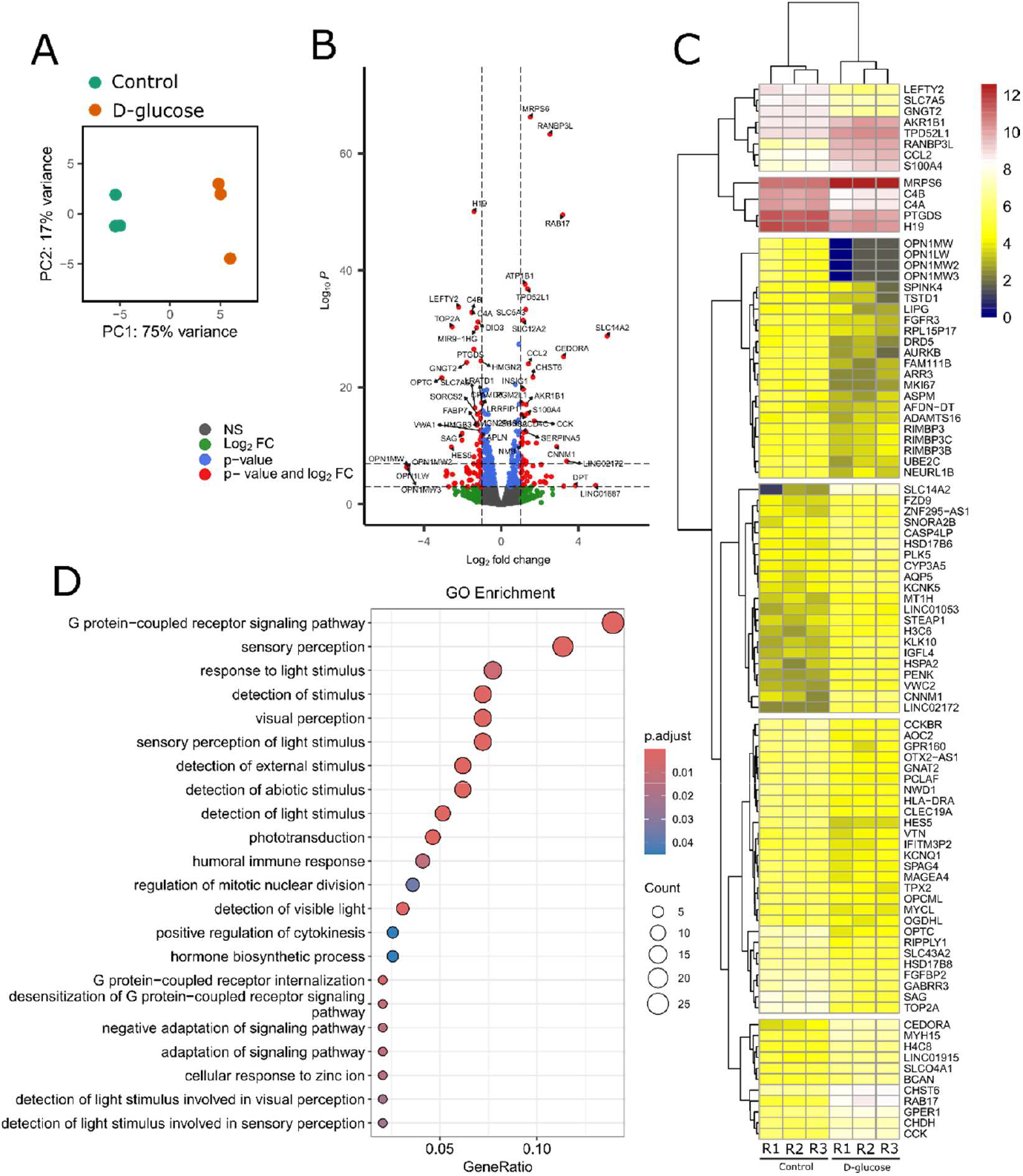
Transcriptomic analysis of retinal organoids exposed to hyperglycaemia. **A)** Principal component analysis (PCA) plot showing the clustering of transcriptomic profiles. Control and D-glucose-treated groups form distinct clusters. **B)** Volcano plot illustrating differential gene expression between retinal organoids treated with D-glucose and controls. Significantly upregulated and downregulated genes are highlighted based on adjusted p-value and log2 fold change. **C)** Heatmap displaying expression patterns of differentially expressed genes. **D)** Gene Ontology (GO) enrichment analysis of all differentially expressed genes (p< 0.05, log2FoldChange >= 1).

We found 251 differentially expressed genes (p < 0.05, log2FoldChange >= 1) (**Supplementary Table S1**). Among the downregulated genes, *OPN1MW, OPN1LW, ARR3* and *SAG* stood out as key photoreceptor-specific genes that exhibited significantly reduced expression under hyperglycaemic conditions (**Fig. 4B, C**). These genes are critical for cone and rod photoreceptor function, and their downregulation highlights the vulnerability of photoreceptor differentiation and maturation to elevated glucose levels. Gene Ontology (GO) enrichment analysis (**Fig. 4D, S2**) revealed that the most significantly enriched pathways were G protein-coupled receptor signalling pathway, sensory perception, response to light stimulus, and visual perception.

To evaluate the consistency and complementarity of our findings, we conducted a comparative analysis between our NGS dataset and the publicly available dataset GSE160306. The GSE160306 dataset consists of transcriptomic profiles from human retinal samples, including control and diabetic retinopathy conditions, enabling a comprehensive exploration of gene expression changes associated with hyperglycaemia and DR ^9^. We identified 50 overlapping differentially expressed genes (p < 0.05) between our dataset and GSE160306 (**Fig. S3A, Supplementary Table S2**). Gene Ontology (GO) analysis of these shared genes revealed that they play role in key biological processes including gliogenesis, eye development, visual and sensory system development (**Fig. S3B**). Next, both datasets were subjected to GO analysis to elucidate enriched biological processes and pathways. Our primary focus was to assess how the dataset GSE160306 integrates into the pathways identified in our GO analysis, providing a comprehensive view of shared and distinct mechanisms. Consistently with our datasets, we found significant overlaps in pathways related to retina including eye development, visual and sensory system development, gliogenesis, and glial cell differentiation (**Fig. S3C**). One of the notable findings from integrating GSE160306 into our dataset was the strong emphasis on photoreceptor-related gene expression changes under hyperglycaemic conditions. This aligned with our observations of altered photoreceptor-related pathways in hyperglycaemic organoids.

### Hyperglycaemia increases oxidative stress response and decreases expression of genes associated with proliferation

To further investigate the molecular impacts of hyperglycaemia on retinal organoids, we analysed gene expression related to oxidative stress, apoptosis, proliferation, and cell-specific markers for various retinal cell types. Genes associated with oxidative stress regulation showed differential expression patterns. Notably, *DUOX1* and *DUOX2* were upregulated in the D-glucose condition (**Fig. 5A, Fig. S4**), indicating an active response to elevated oxidative stress. In contrast, other key oxidative stress-related genes, including *CAT, SOD1, SOD2*, and *NOX1, NOX3*, and *NOX4*, displayed no profound changes. These findings suggest a complex, context-dependent oxidative stress response in retinal organoids under hyperglycaemic conditions, wherein some pathways are activated while others remain unaffected. In the apoptosis-related gene set (**Fig. 5B**), no pro-apoptotic genes were upregulated in response to D-glucose. However, the anti-apoptotic gene *BIRC5* (survivin) was downregulated, indicating a subtle shift in the balance of apoptotic regulation. Despite this, the overall apoptotic gene expression profile reflects a tightly regulated response, potentially preventing an overt induction of apoptosis under hyperglycaemic stress. Proliferation markers (**Fig. 5C**), including *CCND1, MKI67, E2F2, TOP2A*, and *PCNA*, were consistently downregulated in the D-glucose condition, indicating the inhibitory effect of hyperglycaemia on cellular proliferative activity, likely contributing to impaired maturation during chronic glucose exposure.

**Figure 5.**
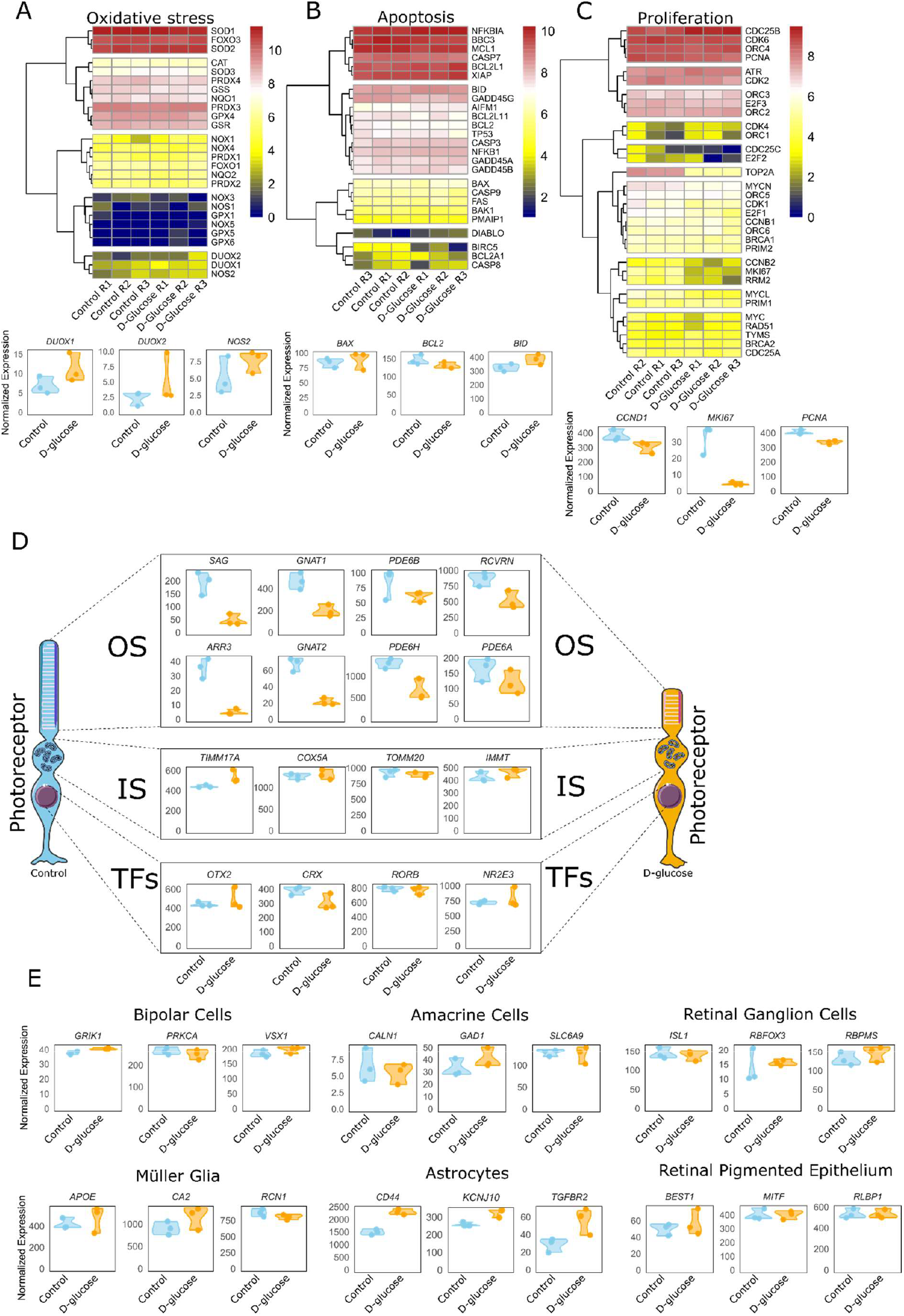
Gene expression analysis of oxidative stress, apoptosis, proliferation, and cell-specific markers in retinal organoids under hyperglycaemic conditions. **A)** Heatmap and violin plots showing the expression patterns of oxidative stress-related genes in retinal organoids treated with D-glucose compared to controls. Key genes include DUOX1, DUOX2, and NOS2. **B)** Heatmap and violin plots for apoptosis-related genes, including BAX, BCL2, and BID. **C)** Heatmap and violin plots for proliferation markers such as CCND1, MKI67, and PCNA. **D)** Expression of photoreceptor-specific genes; outer segment (OS) (rod-specific: SAG, GNAT1, PDE6B, RCVRN; cone-specific: ARR3, GNAT2, PDE6H, PDE6A); inner segment (IS) (TIMM17A, COX5A, TOMM20, IMMT); photoreceptor-specific transcription factors (TFs) (OTX2, CRX, RORB, NR2E3), **E)** Expression of cell-specific markers across different retinal cell types, including bipolar cells (GRIK1, VSX1, PRKCA), amacrine cells (CALN1, GAD1, SLC6A9), retinal ganglion cells (ISL1, RBFOX3, RBPMS), Müller glia (APOE, CA2, RCN1), astrocytes (CD44, KCNJ10, TGFBR2), and retinal pigmented epithelium (BEST1, MITF, RLBP1). Data represent individual data points across three biological replicates for each condition.

Cell-type-specific gene expression analysis (**Fig. 5D**) revealed differential responses among retinal cell markers. Cone and rod photoreceptor OS markers, including *SAG, GNAT1, PDE6B, RCVRN, ARR3, GNAT2, PDE6H*, and *PDE6A*, were markedly downregulated under hyperglycaemic conditions, aligning with previous findings of impaired photoreceptor-specific gene expression. Interestingly, inner segment (IS) markers, such as *TIMM17A, COX5A, TOMM20*, and *IMMT*, along with photoreceptor-specific transcription factors including *OTX2, CRX, NR2E3*, and *RORB*, remained largely unaffected. This suggests that hyperglycaemia primarily disrupts downstream photoreceptor differentiation and functional maturation, rather than early developmental programming. The selective vulnerability of OS-associated genes suggests that photoreceptor outer segments are particularly sensitive to glucose-induced stress, potentially contributing to progressive photoreceptor dysfunction in DR.

In contrast, markers for bipolar cells (*GRIK1, VSX1, PRKCA*), amacrine cells (*CALN1, GAD1, SLC6A9*), retinal ganglion cells (*ISL1, RBFOX3, RBPMS*), Müller glia (*APOE, CA2, RCN1*), and retinal pigment epithelial cells (*BEST1, MITF, RLBP1*) remained largely unchanged. These findings highlight a notable resilience of these retinal cell types to hyperglycaemic stress, suggesting that photoreceptors are particularly vulnerable to glucose-induced damage. Interestingly, astrocyte markers (*CD44, KCNJ10*, and *TGFBR2*) were upregulated under hyperglycaemia. This suggests an increased gliotic response or astrocyte activation, which may represent a compensatory mechanism to counteract hyperglycaemia-induced stress in the retinal environment. This astrocyte activation might also play a role in mediating inflammatory or neuroprotective responses during chronic hyperglycaemia. These findings indicate the cell-type-specific impacts of hyperglycaemia on retinal organoids, with photoreceptors being the most affected, while other retinal cell types exhibit varying degrees of resilience.

## Discussion

In this study, we utilized human retinal organoids as a model to investigate the impact of hyperglycaemia on retinal development across various stages of differentiation. Our results demonstrate that while early-stage retinal organoids exhibit resilience to hyperglycaemic conditions, prolonged exposure at later stages induces significant molecular and cellular disruptions, particularly in photoreceptor differentiation and oxidative stress responses.

In the early stages of differentiation (D30+28 and D90+28), retinal organoids treated with hyperglycaemic conditions showed no significant differences in morphology or structure compared to controls. These findings suggest that the early differentiation processes, such as neural induction and retinal progenitor specification, remain unaffected by hyperglycaemic stress. One possible explanation is that early-stage retinal cells have distinct metabolic demands. Given their high metabolic rate, photoreceptor progenitors and other early-stage retinal cells may possess an intrinsic ability to withstand hyperglycaemic conditions more effectively than their mature counterparts ^10,11^. Supporting this idea, *in vitro* studies have shown that photoreceptor progenitors do not exhibit significant loss under high-glucose exposure, indicating that metabolic requirements could play a key role in their survival ^8^. This metabolic resilience may also indicate that the detrimental effects of hyperglycaemia become more pronounced as organoids undergo further differentiation, paralleling the progressive nature of diabetic retinopathy observed *in vivo*. However, as organoids progressed to later stages of differentiation (D150+28), hyperglycaemic conditions induced significant transcriptional changes, particularly in photoreceptor-specific genes. Genes such as *OPN1SW, OPN1MW, OPN1LW*, and *RHODOPSIN* (**Fig. 2B**), along with additional markers (**Fig. 5D**), were markedly downregulated, highlighting the vulnerability of both cone and rod photoreceptors to prolonged glucose exposure. This progressive impairment mirrors the pathophysiology of DR, where metabolic dysregulation increasingly affects photoreceptors over time ^12^. Interestingly, L-glucose, the osmotic control, also led to a significant downregulation of *OPN1MW*, despite being non-metabolizable. In contrast, all other tested genes remained unaffected under L-glucose conditions, indicating that the effects observed in D-glucose-treated organoids are likely a consequence of hyperglycaemia rather than osmotic stress. The mechanisms underlying this L-glucose-induced downregulation remain unclear but may involve osmotic stress responses, altered nutrient sensing, or unexpected signalling interactions ^13–15^. Despite these L-glucose effects, the stronger and broader transcriptional repression observed in D-glucose-treated organoids supports a hyperglycaemia-specific mechanism affecting photoreceptor maturation and function.

Notably, transcription factors critical for photoreceptor development, such as *CRX* and *NRL*, remained unchanged, suggesting that hyperglycaemia selectively impairs photoreceptor terminal differentiation and function rather than their initial specification. This aligns with the concept that mature photoreceptors, which rely heavily on oxidative metabolism, are particularly susceptible to prolonged metabolic stress, further supporting the parallels between glucose-induced transcriptional changes in organoids and photoreceptor degeneration in diabetic retinopathy ^12^. These findings are consistent with the zebrafish model study by Titialii-Torres et al. ^5^, which demonstrated photoreceptor vulnerability under hyperglycaemic conditions, characterized by shortened OS. While our model similarly revealed significant downregulation of OS-specific photoreceptor genes, the transcriptomic resolution provided by human retinal organoids offered additional insights, highlighting disruptions in phototransduction pathways and the downregulation of genes whose products are critical for OS function and structure. Interestingly, our data sets emphasize oxidative stress as a central driver of hyperglycaemic damage. The observed upregulation of *DUOX1* and *DUOX2* aligns with the findings of increased ROS production ^16,17^, further corroborating the conserved role of oxidative stress in photoreceptor dysfunction. However, other oxidative stress-related genes showed no significant changes, suggesting a nuanced and context-dependent oxidative response. In addition to harmful effects on photoreceptors, we observed a downregulation of genes associated with proliferative activity, including *CCND1, MKI67*, and *PCNA*. This indicates that hyperglycaemic conditions suppress cell proliferation, potentially limiting the growth and maturation of retinal organoids.

Cell-type-specific gene expression analysis revealed differential resilience across retinal cell types. While OS photoreceptor markers were downregulated, markers for bipolar cells, amacrine cells, ganglion cells, Müller glia, and retinal pigment epithelial cells remained largely unaffected. This indicates the cell-type-specific resilience of retinal organoids to hyperglycaemic stress. However, astrocyte markers, including *CD44, GFAP*, and *TGFBR2*, were upregulated, suggesting an activation of glial stress responses. Both our data and the findings of Titialii-Torres et al. ^5^ reported glial activation as a key feature of hyperglycaemia, indicating that reactive gliosis represents a common early response to retinal metabolic stress, contributing to both neuroprotection and inflammation.

Our findings demonstrate that retinal organoids effectively model the effects of hyperglycaemia on retinal development, revealing significant oxidative stress responses, photoreceptor vulnerability, and glial activation. These results align with and expand upon the study by de Lemos et al. ^8^. Consistent with their observations, we detected significant oxidative stress responses in retinal organoids exposed to hyperglycaemia. While their study identified retinal ganglion cells and amacrine cells as early targets of hyperglycaemia, our analysis revealed resilience in markers for these cell types. This could be explained by differences in experimental design, as their study employed much higher concentrations of D-glucose (50 and 75 mM), levels that far exceed physiological conditions and approach cytotoxicity. Such extreme hyperglycaemia likely triggered more severe stress responses, overwhelming cellular defences and leading to widespread damage. This heightened stress response was associated with increased glial reactivity, marked by elevated *GFAP* expression. Our data similarly showed upregulation of *GFAP* and *CD44*, indicating reactive gliosis in response to hyperglycaemia. This glial activation likely represents a common early response, contributing to both neuroprotection and inflammation in the hyperglycaemic retinal environment.

Comparison of our datasets with publicly available datasets ^9^ further validated our findings, despite the inherent biological differences between the models. Our data, derived from retinal organoids, lack key components such as immune cells, vasculature, and systemic influences, which are present in human retinal samples. Nevertheless, both datasets revealed significant overlaps in pathways related to retinal development, gliogenesis, and photoreceptor dysfunction. However, there were some differences in the extent to which certain pathways were enriched. For instance, inflammatory and angiogenic processes were more pronounced in the human retinal dataset, likely due to the presence of vascular and immune components. These differences likely contribute to the distinct pathway representations observed between models. While retinal organoid models effectively capture DR-associated neurodegenerative processes, *in vivo* systems provide essential insights into the vascular and immune components that drive disease progression. Ongoing efforts to generate vascularized retinal organoids or modelling blood-retinal barrier may help bridge this gap, allowing for a more accurate representation of *in vivo* pathophysiological mechanisms and enhancing the translational potential of these models ^18^. Despite this, the model is important as it revealed a neurotoxic effect of hyperglycaemia, highlighting specific pathways and molecular mechanisms underlying diabetic retinopathy that can be further explored in translational research.

Retinal organoids represent a highly effective model for studying the harmful effects of hyperglycaemia on the retina, capturing key features such as oxidative stress responses, photoreceptor vulnerability, and glial activation. Given their close resemblance to human retinal development ^19–21^, they provide a powerful platform for investigating how maternal hyperglycaemia impacts foetal retinal formation and its long-term consequences. Furthermore, this model enables the development and testing of potential therapeutic strategies to mitigate hyperglycaemia-induced retinal damage, including antioxidant treatments, metabolic modulators, and neuroprotective interventions.

## Methods

### Human induced pluripotent stem (hiPS) cell culture

The hiPS cell lines N5 and M8, derived from skin fibroblasts of healthy donors, were used in this study as previously described ^22–24^. The cells were maintained in Essential 8 culture medium (Gibco) supplemented with 1× Penicillin-Streptomycin solution (Biosera) on plates coated with Vitronectin (Thermo Fisher Scientific). hiPS cells were cultured at 37 °C in a humidified incubator with 5% CO2 to ensure optimal growth conditions. For routine passaging, hiPS cells were dissociated using 0.5 mM EDTA, following standard protocols for gentle handling of pluripotent cells. Cell culture media was refreshed daily, and cells were passaged every 4–5 days.

### Retinal organoids differentiation

Retinal organoids were generated following previously reported protocols ^22–25^ with slight modifications. hiPS cells maintained in Essential 8 medium (Gibco) were seeded into U-shaped, cell-repellent 96-well plates (Cellstar) at a density of 5000 cells per well. After 48 hours (designated as day 0 of differentiation), the culture medium was replaced with a growth factor-free chemically defined medium (gfCDM). The gfCDM comprised 45% Iscove’s Modified Dulbecco’s Medium (IMDM, Gibco), 45% Ham’s F12 Nutrient Mix (F12, Gibco), 10% KnockOut Serum Replacement (Gibco), 1% chemically defined lipid concentrate (Gibco), 1% Penicillin-Streptomycin Solution (Biosera), and 10 μM β-mercaptoethanol (Sigma-Aldrich). On day 6, recombinant human BMP4 (Peprotech) was added to the medium at a final concentration of 1.5 nM. Half of the medium volume was replaced with fresh medium every three days. On day 18, the gfCDM was replaced with a neuroretinal (NR) medium composed of DMEM/F12 (Gibco), 1% N-2 supplement (Gibco), 1% GlutaMAX supplement (Gibco), 10% foetal bovine serum (FBS, Biosera), 0.5 mM retinoic acid (Sigma), 0.1 mM taurine (Sigma), and 1% Penicillin-Streptomycin Solution (Biosera). Organoids were maintained in the 96-well plates until day 18, after which they were transferred to 10 cm Petri dishes for continued culture.

### Glucose treatment

Retinal organoids at D30, D90, and D150 were exposed to 3 different conditions. The conditions were 1) Control, containing the maintenance medium with the standard glucose concentration of 17,5 mM, 2) D-glucose, containing maintenance medium supplemented with D-glucose (Sigma-Aldrich) to final concentration of 25 mM of D-glucose, 3) L-glucose, containing the maintenance medium supplemented with L-glucose (Sigma-Aldrich) with the final concentration of 25 mM of glucose. L-glucose was used as the osmotic control to D-glucose. The media were changed twice a week for the period of 4 weeks.

### RNA extraction and reverse transcription quantitative real-time PCR (RT-qPCR)

Retinal organoids were washed twice with PBS and dissociated in RNA Blue Reagent (Top-Bio). Samples were either stored at −80 °C or processed immediately. Organoids were homogenized using an insulin syringe, and total RNA was extracted using the Direct-zol™ RNA MicroPrep Kit (ZYMO RESEARCH, Cat. No.: R2062), following the manufacturer’s protocol. Complementary DNA (cDNA) was synthesized from the extracted RNA using the High-Capacity cDNA Reverse Transcription Kit (Applied Biosystems™). Quantitative PCR was carried out with SYBR Green I (Top-Bio) using the LightCycler® 480 system (Roche). Primer sequences are listed in **Supplementary Table S3**. The expression levels of target genes were normalized against the housekeeping gene GAPDH, and relative fold changes were calculated using the ΔΔCt method.

### Immunofluorescence staining

Organoids were washed with PBS and fixed in 4% paraformaldehyde (PFA) solution in PBS for 30 minutes at room temperature. After fixation, the PFA was removed, and organoids were washed three times with PBS. They were then incubated overnight in 30% sucrose (Sigma-Aldrich) solution at 4 °C for cryoprotection. Subsequently, organoids were transferred into plastic disposable base molds, the remaining sucrose solution was carefully removed, and the molds were filled with Tissue-Tek® O.C.T. Compound medium (Sakura). The molds containing organoids were placed on dry ice for 15 minutes to solidify the medium and stored at −20 °C or processed immediately. Frozen blocks were sectioned into 7 µm-thick slices using a Leica CM1850 Cryostat (Leica). Slides were allowed to dry at room temperature for 20 minutes before further processing. Sections were washed three times with PBS and incubated for 1 hour in blocking buffer (0.3% Triton-X-100, 5% normal goat serum, in PBS) to reduce nonspecific binding. The blocking buffer was then removed, and the slides were incubated overnight at 4 °C in a humidified chamber with primary antibodies diluted in antibody diluent (0.3% Triton-X-100, 1% BSA, in PBS). Primary antibodies and their concentrations are listed in **Supplementary Table S4**. After incubation with primary antibodies, slides were washed three times with antibody diluent (3 minutes per wash) and then incubated with secondary antibodies for 1 hour at room temperature in a humidified chamber. Secondary antibodies included Goat anti-Rabbit IgG (H+L) Alexa594 (1:1000, ThermoFisher) and Goat anti-Mouse IgG (H+L) Alexa488 (1:1000, ThermoFisher). Cell nuclei were counterstained with DAPI (4′,6-diamidino-2-phenylindole; 1 µl in 1 ml of antibody diluent) for 5 minutes. Slides were then washed five times with PBS followed by one final rinse in distilled water. The sections were mounted using Fluoroshield™ mounting medium (Sigma-Aldrich), and coverslips were sealed with transparent nail polish to prevent movement or drying. Samples were imaged using a Zeiss LSM 880 laser scanning confocal microscope equipped with the AiryscanFast module. Image processing and analysis were performed using the Fiji platform ^26^.

### Human apoptosis array

Organoids were washed thoroughly with PBS (10 washes) to remove residual media and lysed in lysis buffer containing 1% SDS, 10% glycerol, and 50 mM TRIS-HCl (pH 6.8). The lysates were sonicated using an ultrasonic homogenizer (Bandelin Sonopuls HD 2200) to ensure thorough cell disruption and protein solubilization. Protein concentrations in the lysates were quantified and normalized to ensure equal protein loading across samples. The samples were then applied to the Proteome Profiler™ Human Apoptosis Array Kit (R&D Systems, Cat. No. ARY009), following the manufacturer’s protocol. After completing the assay, membranes were visualized using the ChemiDoc™ Touch Imaging System (Bio-Rad). The resulting signals were analysed using the Fiji (ImageJ) software’s Protein Array Analyzer plugin, enabling quantitative evaluation of apoptosis-related protein expression.

### Next Generation Sequencing (NGS)

RNA integrity was checked on the Fragment Analyzer using RNA Kit 15 nt (Agilent Technologies). 500ng of total RNA was used as input for library preparation using QuantSeq 3′ mRNA-Seq FWD with UDI 12 nt Kit (v.2) (Lexogen) in combination with UMI Second Strand Synthesis Module for QuantSeq FWD. Quality control for library quantity and size distribution was done using QuantiFluor dsDNA System (Promega) and High Sensitivity NGS Fragment Analysis Kit (Agilent Technologies). Final library pool was sequenced on NextSeq using High Output Kit v2.5 75 cycles (Illumina) in single-end mode, resulting in a minimum of 10 million reads per sample. Quality check of raw single-end fastq reads was carried out by FastQC ^27^. The adapters and quality trimming of raw fastq reads was performed using Trimmomatic v0.39 ^28^ with settings CROP:250 LEADING:3 TRAILING:3 SLIDINGWINDOW:4:5 MINLEN:35. Trimmed RNA-Seq reads were mapped against the mouse genome (hs38) and Ensembl GRCh38-p10 annotation using STAR v2.7.3a ^29^ as splice-aware short read aligner and default parameters except -- outFilterMismatchNoverLmax 0.66 and -- twopassMode Basic. Quality control after alignment concerning the number and percentage of uniquely- and multi-mapped reads, rRNA contamination, mapped regions, read coverage distribution, strand specificity, gene biotypes and PCR duplication was performed using several tools namely RSeQC v4.0.0 ^30^, Picard toolkit v2.25.6 ^31^, Qualimap v.2.2.2 ^32^. NGS datasets were processed in RStudio using packages: DESeq2 ^33^, biomaRt ^34^, Rsubread ^35^, EnhancedVolcano ^36^, pheatmap ^37^, and clusterProfiler ^38^. NGS data are accessible at GEO database: GSE290024.

### Data analysis and statistics

Statistical analysis and data visualization were performed using RStudio, employing libraries: ggstatsplot ^39^ and ggplot2 ^40^. All experiments included at least three independent biological replicates. For RT-qPCR, each biological replicate was analysed in technical triplicates, with individual data points displayed in the corresponding graphs. Statistical significance was assessed using a two-tailed Student’s t-test, and p-values are reported in each panel. Comparisons with p < 0.05 were considered statistically significant.

## Acknowledgement

This study was supported by the funds by the Faculty of Medicine MU (MUNI/A/1738/2024 and MUNI/LF-SUp/1197/2022) and Macular Society/Diabetes UK 23/0006592, MRC - APP13221. We acknowledge the core facility CELLIM supported by the Czech Bioimaging large RI project (LM2023050 funded by MEYS CR) for their support with obtaining scientific data presented in this presentation. We acknowledge the CF Genomics and the CF Bioinformatics supported by the NCMG research infrastructure (LM2023067 funded by MEYS CR) for their support with obtaining scientific data presented in this paper. The CREATIC project is funded by the European Union (Grant Agreement No. 101059788).

## Data availability

The transcriptomic data set is available through GEO under accession number GSE290024. Any additional requests for information can be directed to, and will be fulfilled by, the corresponding author.

## Supplementary figures

**Figure S1:**
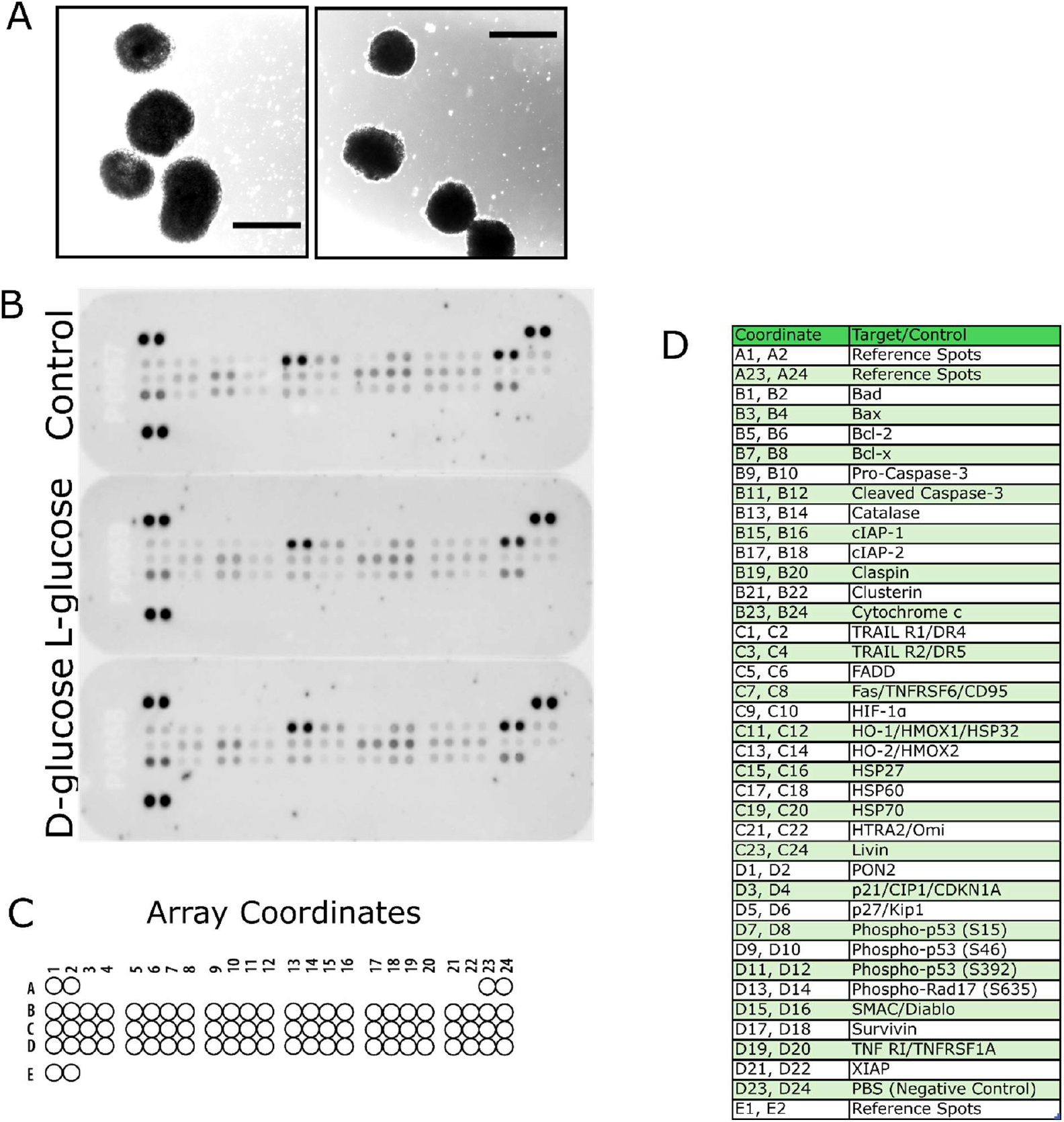
**A)** Images of retinal organoids at D30 cultured for additional 7 days in 5.5 mM D-glucose. We observed increased cell death and organoid degradation. Scale bars represents 500 µm. **B)** Images of protein arrays used to analyse apoptosis-related markers in retinal organoids exposed to hyperglycaemic conditions. C) Corresponding coordinates on the chip indicating the placement of individual protein spots. D) A reference table listing the coordinates and the respective protein symbols for the detected apoptosis-related markers.

**Figure S2:**
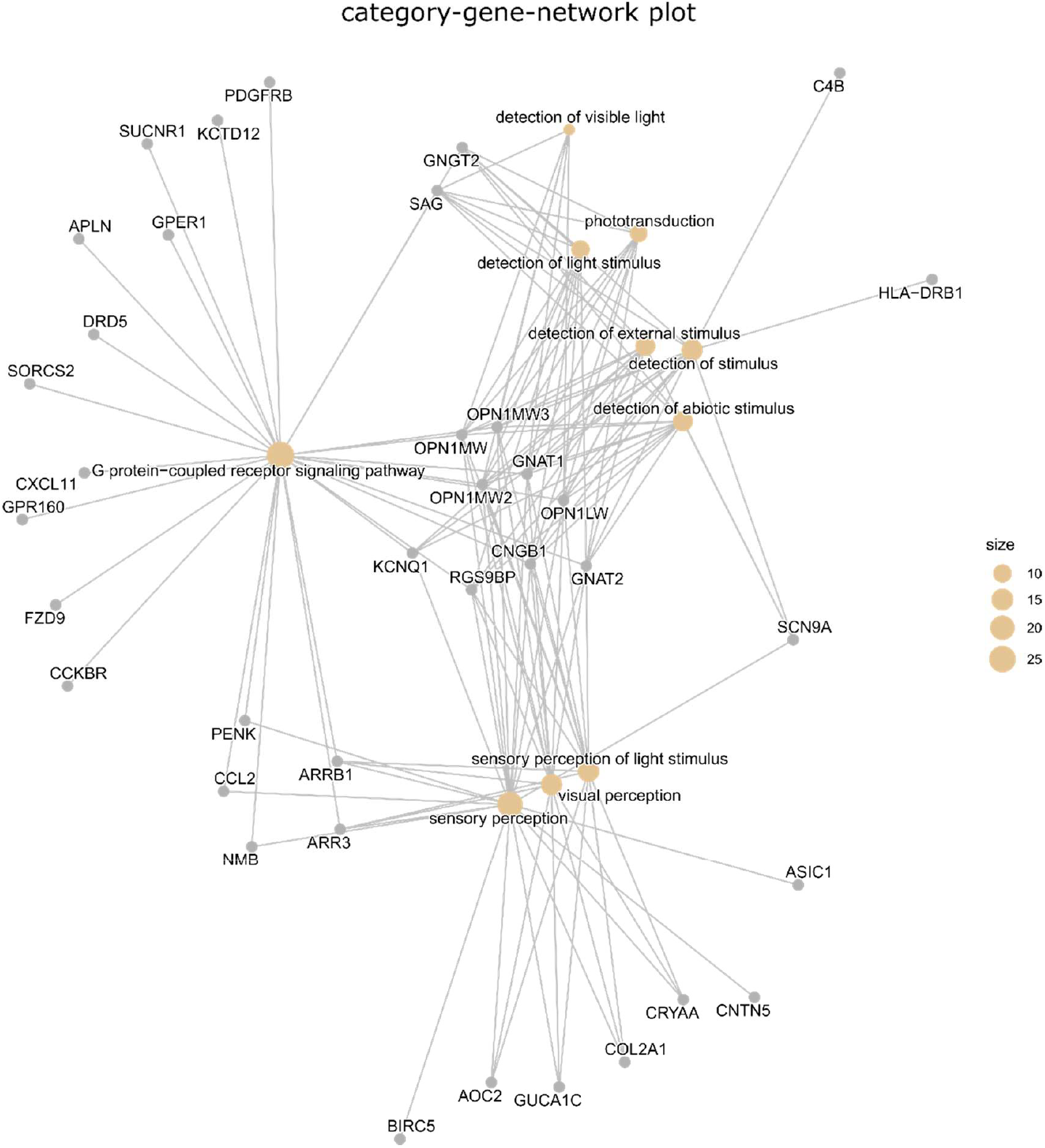
Category-gene network plot of GO enrichment analysis (Top 10 categories are shown)

**Figure S3:**
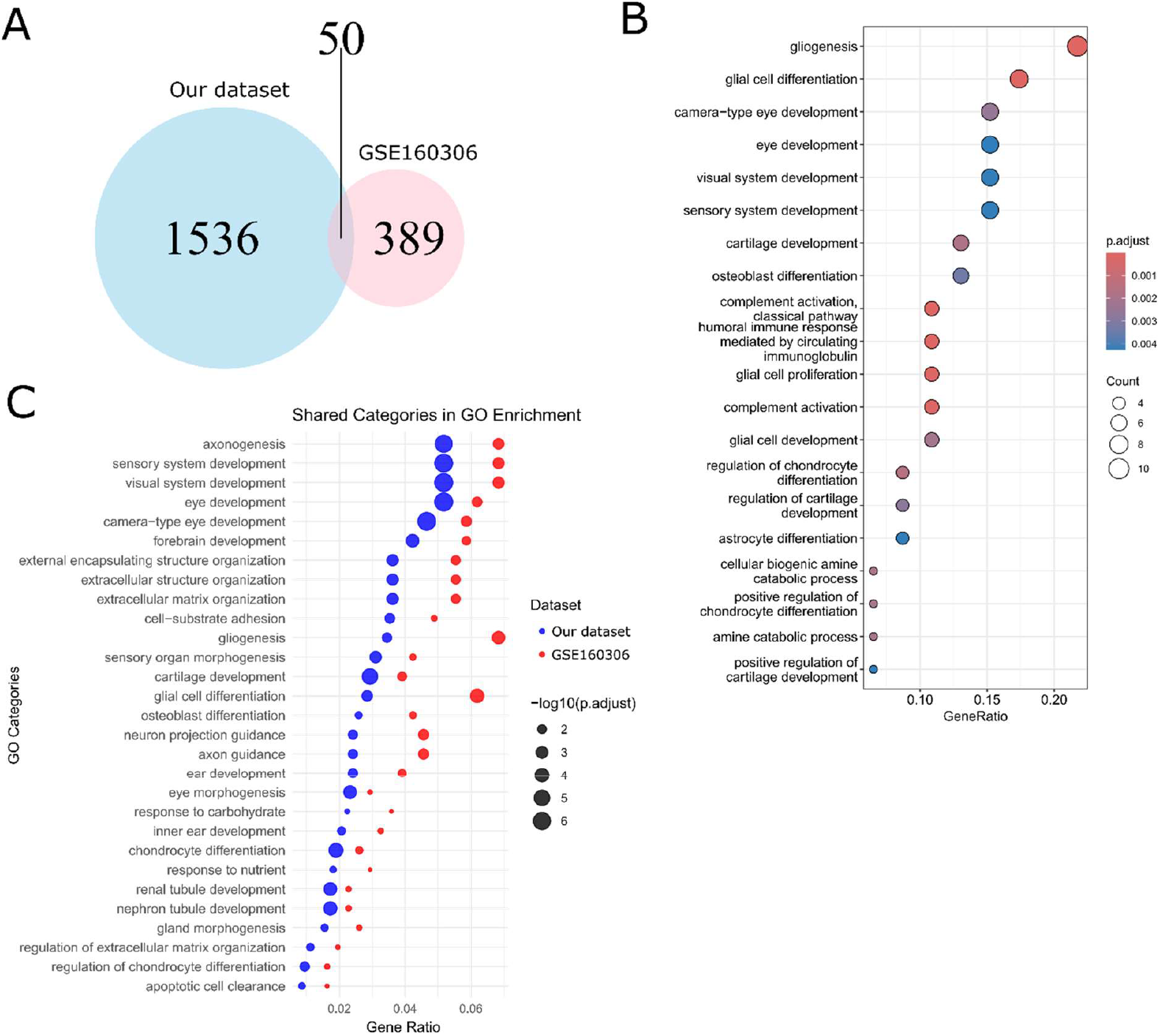
**A)** Venn diagram depicting the overlap of differentially expressed genes (p< 0.05) identified in hyperglycaemia-exposed retinal organoids (blue) and a publicly available diabetic retinopathy dataset GSE160306 (pink). The shared genes (intersection) represent common genes affected in both datasets. **B)** Dot plot showing enriched Gene Ontology (GO) terms for the shared genes. **C)** Dot plot representing enriched shared pathways in both dataset (blue dots represent our dataset, while red dots represent GSE160306).

**Figure S4:**
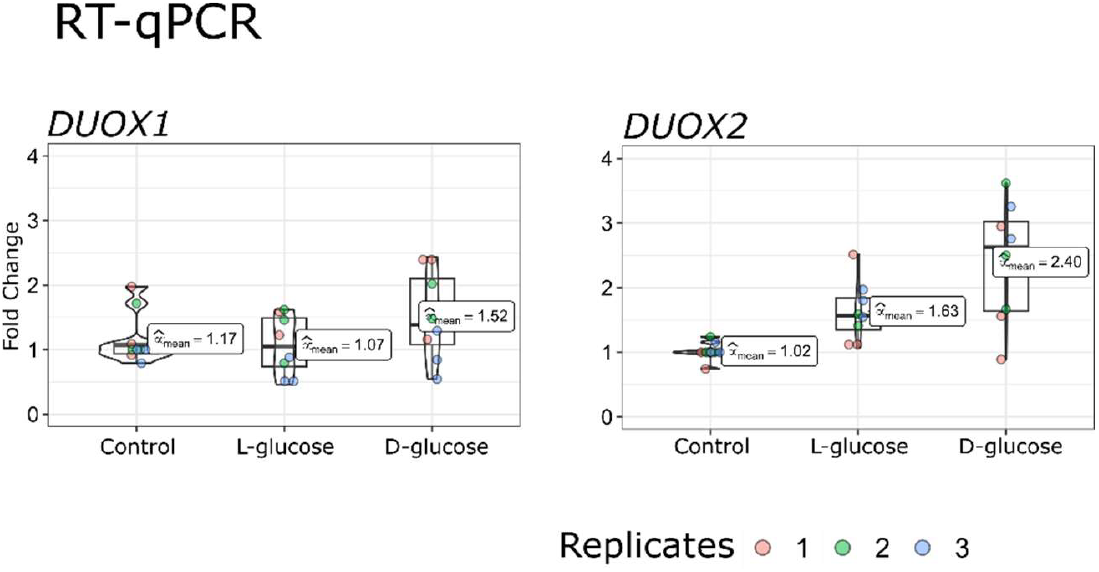
RT-qPCR analysis of DUOX1 and DUOX2 expression in late stage (D150+28) retinal organoids treated with control or L-/D-glucose.

